# Resolving the complex *Bordetella pertussis* genome using barcoded nanopore sequencing

**DOI:** 10.1101/381640

**Authors:** Natalie Ring, Jonathan Abrahams, Miten Jain, Hugh Olsen, Andrew Preston, Stefan Bagby

**Affiliations:** Department of Biology and Biochemistry, University of Bath, Bath, UK; Nanopore Group, University of California Santa Cruz, California, USA

## Abstract

The genome of *Bordetella pertussis* is complex, with high GC content and many repeats, each longer than 1,000 bp. Short-read DNA sequencing is unable to resolve the structure of the genome; however, long-read sequencing offers the opportunity to produce single-contig *B. pertussis* assemblies using sequencing reads which are longer than the repetitive sections. We used an R9.4 MinION flow cell and barcoding to sequence five *B. pertussis* strains in a single sequencing run. We then trialled combinations of the many nanopore-user-community-built long-read analysis tools to establish the current optimal assembly pipeline for *B. pertussis* genome sequences. Our best long-read-only assemblies were produced by Canu read correction followed by assembly with Flye and polishing with Nanopolish, whilst the best hybrids (using nanopore and Illumina reads together) were produced by Canu correction followed by Unicycler. This pipeline produced closed genome sequences for four strains, revealing inter-strain genomic rearrangement. However, read mapping to the Tohama I reference genome suggests that the remaining strain contains an ultra-long duplicated region (over 100 kbp), which was not resolved by our pipeline. We have therefore demonstrated the ability to resolve the structure of several *B. pertussis* strains per single barcoded nanopore flow cell, but the genomes with highest complexity (e.g. very large duplicated regions) remain only partially resolved using the standard library preparation and will require an alternative library preparation method. For full strain characterisation, we recommend hybrid assembly of long and short reads together; for comparison of genome arrangement, assembly using long reads alone is sufficient.

**DATA SUMMARY:**

1. Final sequence read files (fastq) for all 5 strains have been deposited in the SRA, BioProject PRJNA478201, accession numbers SAMN09500966, SAMN09500967, SAMN09500968, SAMN09500969, SAMN09500970
2. A full list of accession numbers for Illumina sequence reads is available in Table S1
3. Assembly tests, basecalled read sets and reference materials are available from figshare: https://figshare.com/projects/Resolving_the_complex_Bordetella_pertussis_genome_using_barcoded_nanopore_sequencing/31313
4. Genome sequences for *B. pertussis* strains UK36, UK38, UK39, UK48 and UK76 have been deposited in GenBank; accession numbers: CP031289, CP031112, CP031113, QRAX00000000, CP031114
5. Source code and full commands used are available from Github: https://github.com/nataliering/Resolving-the-complex-Bordetella-pertussis-genome-using-barcoded-nanopore-sequencing

**IMPACT STATEMENT:** Over the past two decades, whole genome sequencing has allowed us to understand microbial pathogenicity and evolution on an unprecedented level. However, repetitive regions, like those found throughout the *B. pertussis* genome, have confounded our ability to resolve complex genomes using short-read sequencing technologies alone. To produce closed *B. pertussis* genome sequences it is necessary to use a sequencing technology which can generate reads longer than these problematic genomic regions. Using barcoded nanopore sequencing, we show that multiple *B. pertussis* genomes can be resolved per flow cell. Use of our assembly pipeline to resolve further *B. pertussis* genomes will advance understanding of how genome-level differences affect the phenotypes of strains which appear monomorphic at nucleotide-level.

This work expands the recently emergent theme that even the most complex genomes can be resolved with sufficiently long sequencing reads. Additionally, we utilise a more widely accessible alternative sequencing platform to the Pacific Biosciences platform already used by large research centres such as the CDC. Our optimisation process, moreover, shows that the analysis tools favoured by the sequencing community do not necessarily produce the most accurate assemblies for all organisms; pipeline optimisation may therefore be beneficial in studies of unusually complex genomes.

## INTRODUCTION

*Bordetella pertussis* is the pathogenic bacterium which causes most cases of whooping cough (pertussis). Pertussis was a greater medical burden prior to the international introduction of vaccination in the 1940s and 1950s. Widespread vaccine uptake greatly reduced incidence of the disease in developed countries. Original whole-cell vaccines were replaced by new acellular vaccines throughout the 1990s and early 2000s. The acellular vaccines contain three to five *B. pertussis* protein antigens. All versions contain pertactin (Prn), pertussis toxin (Pt) and filamentous haemagglutinin (FHA), and some also contain one or both of the fimbrial proteins Fim2 and Fim3. Despite continued high levels of coverage of pertussis vaccination, since the early 1990s the number of cases of whooping cough has increased in many countries [1, 2].

Suggested causes for this resurgence include improved diagnostic tests and awareness, waning immunity as a result of the switch to acellular vaccination, and genetic divergence of circulating *B. pertussis* from the vaccine strains due to vaccination-induced selection pressure [3-5]. A global survey of strains from the pre-vaccine, whole cell vaccine and acellular vaccine eras showed that the genome of *B. pertussis*, a traditionally monomorphic and slowly-evolving organism, has been evolving since the introduction of the whole-cell vaccine [6]. Analysis of strains from several recent epidemics showed that this evolution has been particularly rapid in the genes encoding vaccine antigens since the switch to the acellular vaccine [7-10].

The *B. pertussis* genome contains many repeats of an insertion sequence (IS), IS481. Recombination has led to the appearance of up to 300 copies of IS481, which is 1,053 bp long. A smaller number of copies of IS1002 (1,040 bp) and IS1663 (1,014 bp) contribute further complexity to the genome. These regions of repetition mean that assembly of closed, single-contig *B. pertussis* genomes using short-read sequencing, which produces reads shorter than the IS repeats, has been particularly difficult: most genome sequences available on NCBI comprise several hundred contigs, or at least one contig per IS element copy. Over the last decade, many studies have shown that reads longer than the longest repeat are required to resolve regions of high complexity [11-18].

In 2016, Bowden *et al.* [19] were the first to use long reads, together with Illumina short reads, to conduct a survey of *B. pertussis* strains which had circulated during two whooping cough epidemics, in the US, in 2010 and 2012. Assembling closed genomes for these epidemic isolates revealed extensive genomic arrangement differences between isolates which appeared to be otherwise closely related. They concluded that further comprehensive whole genome studies are required to fully understand the international resurgence of whooping cough. More recently, Weigand *et al.* showed that the *B. pertussis* genome has undergone, and continues to undergo, structural rearrangement on a relatively frequent basis [20].

Bowden *et al.* and Weigand *et al.* both used Pacific Biosciences (PacBio) long read sequencing, which has high start-up costs, and lacks the portability needed for on-the-ground epidemic surveillance. In contrast, Oxford Nanopore Technology (ONT)’s MinION nanopore sequencer has nominal start-up cost. Recent improvements to flow cell yield and the introduction of barcoded library preparation make per-sample MinION costs comparable to those of PacBio or Illumina [15, 21, 22]. In addition, the pocket-sized MinION is portable, enabling in-the-field sequencing [23-25].

Here we test the ability of barcoded nanopore sequencing to resolve the genomes of five *B. pertussis* strains from a UK epidemic, which were previously unclosed and comprised many contigs assembled using short reads sequenced with Illumina’s MiSeq [7]. We subsequently trial a variety of available data analysis tools to develop a bioinformatics pipeline capable of rapid assembly of accurate, closed *B. pertussis* genome sequences using only long reads or, if available, short reads together with long reads.

## METHODS

Full method and bioinformatics procedures are described at: https://github.com/nataliering/Resolving-the-complex-Bordetella-pertussis-genome-using-barcoded-nanopore-sequencing

All data analysis was carried out using the Medical Research Council’s Cloud Infrastructure for Microbial Bioinformatics (CLIMB) [26].

### Strain isolation and Illumina sequencing

Five strains originally isolated during the UK 2012 whooping cough epidemic were obtained from the National Reference Laboratory, Respiratory and Vaccine Preventable Bacteria Reference Unit, at Public Health England. Short-read sequencing data were generated previously, using genomic DNA (gDNA) extracted using DNeasy Blood and Tissue kit (Qiagen), multiplex library preparation and Illumina sequencing [7]. Full details, including accession numbers, are included in **Table S1**.

### DNA extraction

Strains obtained from the National Reference Laboratory were stored at -80°C in phosphate buffered saline (PBS)/20 % glycerol at the University of Bath. These were then grown on charcoal agar plates (Oxoid) for 72 hours at 37°C. All cells were harvested from each plate and resuspended in 3 ml PBS. The optical density of each cell suspension was measured at 600 nm, and volumes of suspension equating to 1.0 OD (~2x10^9^ *B. pertussis* cells) in 180 µl were pelleted in a microcentrifuge for 2 minutes at 12,000 *g*. gDNA was extracted from each pellet using GenElute bacterial genomic DNA kit (Sigma Aldrich) according to manufacturer’s instructions, including the optional RNAase A step and a two-step elution into 200 µl elution buffer (10 mM Tris-HCl, 0.5 mM EDTA, pH 9.0). QuBit fluorometry (dsDNA HS kit, Invitrogen) was used to measure gDNA concentration, and Nanodrop spectrometry (ThermoFisher Scientific) was used to assess gDNA purity.

### Nanopore library preparation and MinION sequencing

1.5 µg of gDNA per strain was concentrated using a 2.5x SPRI clean-up (AMPure XP beads, Beckman Coulter), eluting into 50 µl of nuclease-free (NF) water (Ambion). 2 µl of each sample was used for Bioanalyzer fragment analysis (DNA 12000 kit, Agilent), according to manufacturer’s instructions. The remaining 48 µl was sheared to 20 kb using g-tubes (Covaris), according to manufacturer’s instructions.

To determine whether FFPE (Formalin-Fixed, Paraffin-Embedded) repair improved sequencing yield, half of each sheared sample was end-repaired in a total reaction volume of 62 μl, containing 45 μl gDNA, 6.5 μl NEBNext FFPE repair buffer (New England Biolabs), 2 μl NEBNext FFPE repair mix (NEB) and 8.5 μl NF water. The mixture was incubated at 20°C for 15 minutes. A 1x SPRI clean-up was carried out on the end-repaired samples, eluting into 46 *μ*l NF water.

Sequencing libraries were prepared for all FFPE-repaired and non-FFPE-repaired samples using Oxford Nanopore Technology’s (ONT, Oxford, UK) 1D ligation sequencing kit (SQK-LSK108) with native barcoding (EXP-NBD103), according to manufacturer’s instructions. Ten barcodes were used: one for FFPE-repaired samples from each strain, and one for non-FFPE-repaired sample from each strain. The starting mass of gDNA for each sample was 1.35 μg. After library preparation, different volumes of samples were combined to produce an equi-mass pool for eight samples; two samples had much lower concentration after library preparation so were pooled in full. A total mass of 712.5 ng was pooled in 208.9 μl NF water, which was concentrated to 50 μl by 2.5x SPRI clean-up. Full details of mass pooled per sample are given in **Table S1**. This pooled 50 μl library was used for sequencing adapter ligation.

The final sequencing library was loaded onto an R9.4 flow cell and sequenced for 48 hours using a MinION MK1b device with MinKNOW sequencing software (protocol NC_48h_Sequencing_Run_FLOMIN106_SQK-LSK108). MinKNOW performed concurrent basecalling, outputting sequenced reads in fast5 and fastq format.

### Additional basecalling and demultiplexing

The MinKNOW-basecalled reads were demultiplexed using Porechop (v0.2.1, https://github.com/rrwick/Porechop), which also trimmed adapter sequences. To compare basecaller accuracy, the fast5s were re-basecalled using ONT’s Albacore (v2.1.3), with barcode binning. As suggested in Wick *et al.*’s 2017 protocol [15], Porechop was then used to demultiplex the Albacore reads, keeping only those for which Albacore and Porechop agreed on the bin. This produced three sets, each with 10 bins: 10 barcode bins for MinKNOW + Porechop, 10 barcode bins for Albacore alone, and 10 barcode bins for Albacore + Porechop. The Albacore + Porechop fastqs were deposited in the SRA with accession codes SAMN09500966 to SAMN0900970. Full details of all three read sets are given in **Table S1**.

### Assembly of short-read-only drafts

Assuming the available Illumina data to have typically low error, short-read-only genome sequences were assembled for each strain using ABySS (v2.0.3) [27]. Prior to assembly, reads were prepared using Trimmomatic (v0.34) [28], which trimmed the first 10 bases of each read, and discarded any reads whose five-base sliding-window q-score fell below 32. These assemblies had low contiguity, but theoretically high accuracy.

### Comparison of raw reads

A shell script, “summary_stats”, was used to give total number of reads, mass sequenced and mean read length for each set of raw reads. Summary_stats uses seq_length.py [29] and all_stats. All are available from https://github.com/nataliering/Resolving-the-complex-Bordetella-pertussis-genome-using-barcoded-nanopore-sequencing. Raw % identity was estimated by comparing each read set to the *B. pertussis* reference genome (Tohama I, NC_002929.2). As the UK 2012 strains were not expected to be identical to the 2003 Tohama I sequence, read error was also estimated by comparison with the respective Illumina-only assemblies. The comparison was conducted using BWA MEM [30] and samtools stats [31], which produces a long output file including “error rate” (% identity was calculated from this: 100-(error rate*100)). Raw_error (https://github.com/nataliering/Resolving-the-complex-Bordetella-pertussis-genome-using-barcoded-nanopore-sequencing/blob/master/raw_error) produces a.stats file using this method, given a read set and reference genome. Using the same BWA MEM output, raw read coverage of the Tohama I reference genome was checked using samtools depth.

Finally, raw GC % content was calculated using GC_calculator which outputs the % GC content of a given fasta file (https://github.com/nataliering/Resolving-the-complex-Bordetella-pertussis-genome-using-barcoded-nanopore-sequencing/blob/master/GC_calculator).

These raw statistics were used to determine which set of reads to carry forward to assembler testing (MinKNOW, Albacore or Albacore+Porechop), and whether to use the end-repaired sample or non-end-repaired sample for each strain (or to pool the reads from both).

### Assembly tool testing – nanopore only

The read sets for one barcoded strain, UK36, were used to test a variety of *de novo* assembly strategies. For this, the Albacore+Porechop reads from the FFPE-repaired and non-FFPE-repaired samples were pooled. Four community-built assembly tools were trialled: ABruijn (now called Flye, v1.0 and v2.3.2 respectively), Canu (v1.5), Miniasm with Minimap/Minimap2 (v0.2-r128, v0.2-r123 and v2.0-r299-dirty, respectively) and Unicycler (v0.4.4) [32-35].

Canu has a standalone option to conduct pre-assembly read correction. This was used to correct the 359x coverage UK36 read set to 40x coverage of more accurate reads. Each assembly tool was then trialled with and without pre-assembly read correction. As Canu’s read correction step is relatively CPU time-consuming, an alternative was also trialled. Filtlong (https://github.com/rrwick/Filtlong) does not correct reads, but produces read sets comprising the longest and most accurate reads, up to a given level of coverage; 40x and 100x coverage were trialled here.

Finally, Racon (v.1.2.0) [36] was tested to determine whether the draft assemblies could be improved by post-assembly polishing. After each Racon polish, the accuracy of the assembly produced was estimated. If an improvement was observed, another round of polishing was conducted, up to a total of five rounds. Once two successive rounds of polishing showed no further improvement, no further Racon polishes were conducted. For Unicycler, no manual Racon polishes were conducted, because Racon polishing is part of the Unicycler assembly process. After Racon polishing, each assembly was further polished with a single round of Nanopolish (v0.9.0) [14].

Testing exhaustive combinations of each of these steps produced 28 draft assemblies for each of the assembly tools (see **Table S2** for all combinations).

### Assembly tool testing - hybrid

As Illumina reads were already available for the strains sequenced here, a variety of hybrid *de novo* assembly strategies were also tested. Using Pilon (v1.22) [37], the best nanopore-only assembly for each of the assembly tools was polished with the Illumina reads, up to a total of five rounds. In addition, a hybrid assembly was produced using Unicycler’s hybrid mode, which combines both read sets for assembly, and conducts several rounds of Racon and Pilon polishes automatically. Finally, the hybrid assembly mode of SPAdes (v3.12.0) [38] was tested. These hybrid tests produced another 22 draft assemblies (**Table S2**).

### Assessing assembly accuracy

Summary_stats was used to determine number of contigs, and contig length for each draft assembly. % identity of each draft compared to the Illumina-only draft was estimated using a method developed by Wick *et al.* [39]. Their chop_up_assembly.py and read_length_identity.py scripts were used to generate % identity figures for 10 kbp sections along the entirety of each assembly, and a custom shell script, assembly_identity (https://github.com/nataliering/Resolving-the-complex-Bordetella-pertussis-genome-using-barcoded-nanopore-sequencing/blob/master/assembly_identity) was used to calculate the mean % identity of the whole.

Quality metrics for each assembly were produced using Quast (v4.5) [40] and BUSCO (v1.22) [41]. In addition, a method developed by Watson [42], Ideel (https://github.com/mw55309/ideel) was used to assess the effect of any erroneous indels in the final UK36 hybrid assembly. Ideel was altered slightly: Prokka was used to predict proteins instead of Prodigal, because Prokka can use a set of reference proteins during annotation.

### Comparing genome arrangement

After the best nanopore-only and hybrid assembly pipelines were identified for UK36, the pipelines were used to produce draft assemblies for the remaining four strains. The hybrid assembly for each strain was annotated using Prokka (v1.12) [43], and the genomes were submitted to GenBank (accession numbers CP031289, CP031112, CP031113, QRAX00000000, CP031114).

The arrangement of each nanopore-only assembly was compared to that of each hybrid using progressiveMauve (v20150226 build 10) [44]. Finally, the nanopore-only assemblies for each strain were compared to each other, also using progressiveMauve. Prior to these alignments, each draft was manually rearranged so that the first gene after the *B. pertussis* origin of replication, gidA, was at the beginning of the sequence. gidA_blast (https://github.com/nataliering/Resolving-the-complex-Bordetella-pertussis-genome-using-barcoded-nanopore-sequencing/blob/master/gidA_blast) locates the gidA sequence in the draft to enable manual rearrangement. Later, this same process was used to identify IS element copies in the assembled genomes. If a tool assembled the complementary strand instead of the template (as identified by the results of gidA_blast), a reverse complement of the draft sequence was generated using reverse_complement (https://github.com/nataliering/Resolving-the-complex-Bordetella-pertussis-genome-using-barcoded-nanopore-sequencing/blob/master/reverse_complement).

## RESULTS

### Basecaller comparison

The fast5 sequencing files were basecalled with two different tools: MinKNOW (vJune2017) and Albacore (v2.1.3). The MinKNOW fastq reads were demultiplexed using Porechop (v0.2.1), whilst Albacore performs its own demultiplexing. Due to testing FFPE-repair vs non-FFPE-repair, two read sets existed for each strain sequenced; both sets were pooled and compared to an Illumina-only assembly for each strain to estimate % identity of the raw reads (**Fig. 1a**). The mean identity for the MinKNOW reads was 2.46 % lower than that of the Albacore reads, meaning Albacore produced significantly more accurate basecalls than MinKNOW for our five strains (n=5, paired t-test p<0.001).

**Fig. 1:**
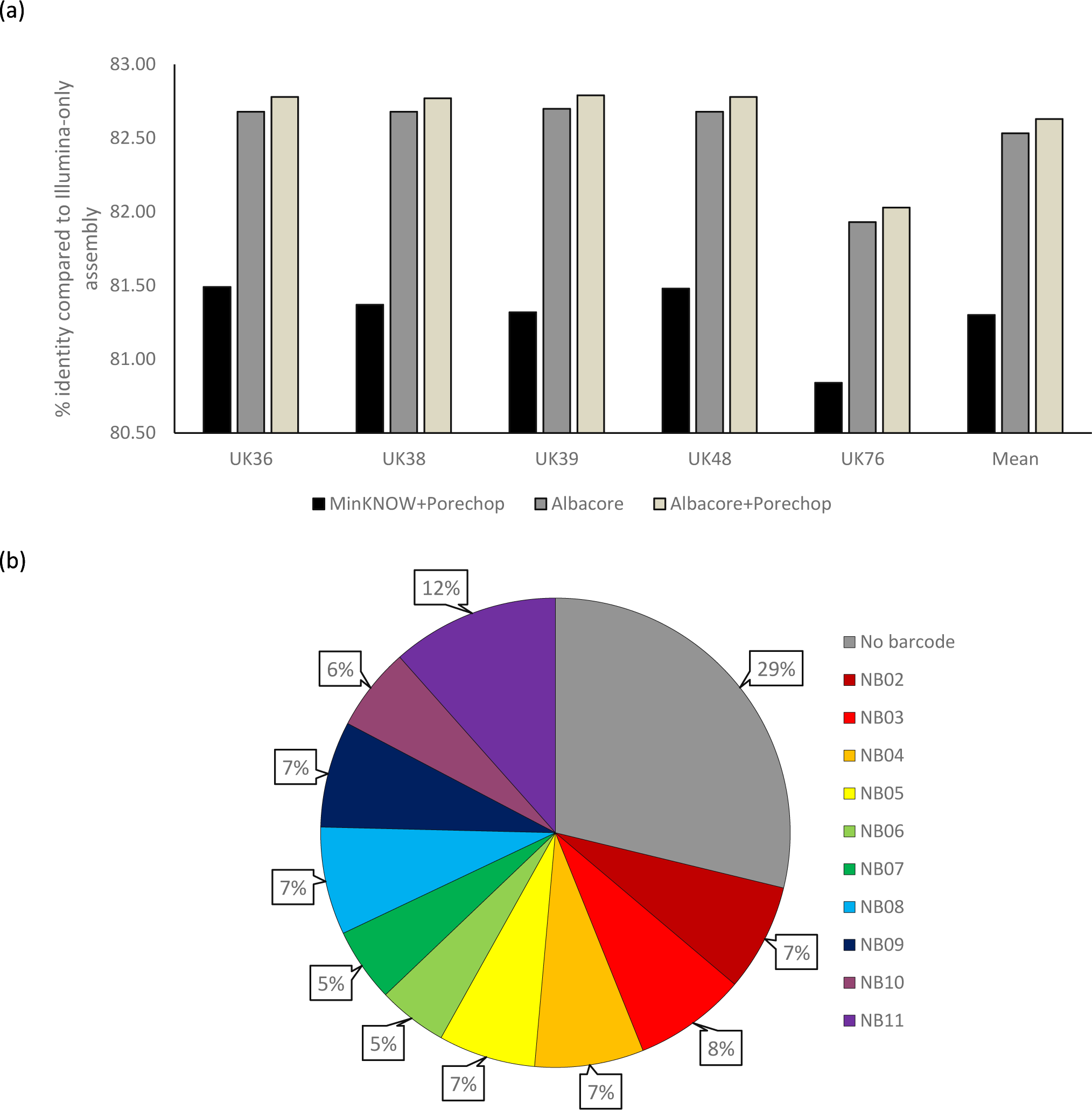
Comparison of basecaller accuracy and barcode distribution. Reads were basecalled and demultiplexed by MinKNOW+Porechop, Albacore or Albacore+Porechop, and compared to an Illumina-only assembly for each strain. As shown in a), both Albacore read sets were more accurate for all five strains, and an additional demultiplexing step with Porechop after Albacore demultiplexing added a mean 0.1 % accuracy. The Albacore+Porechop reads were then used to assess barcode distribution, as shown in b). This showed that a large portion of the raw reads were placed into the “no barcode” bin, meaning Albacore and Porechop either did not agree on a barcode, or no recognisable barcode was present. Otherwise, the barcodes were largely well distributed.

An additional step can also produce greater raw accuracy: the Albacore-demultiplexed reads were re-demultiplexed using Porechop, which keeps only the reads for which both tools agree on the barcode identified. This additional step resulted in a further small but significant improvement in identity compared to Illumina-only assembly: 82.43 % to 82.52 % (n=5, paired t-test p<0.001. Consequently, the reads used for data pipeline development were those that had been basecalled and demultiplexed by Albacore, followed by Porechop re-demultiplexing. For full statistics, see **Table S1**.

### FFPE repair test

Half of each strain’s gDNA sample was repaired with NEBNext FFPE reagents (NEB) to test whether sequencing yield or raw accuracy could be improved by including this optional step prior to library preparation. However, neither the yield (n=5, paired t-test p=0.39) nor the raw accuracy (n=5, paired t-test p=0.937) was significantly improved for the five strains tested. The “FFPE-repaired” and “non-FFPE-repaired” reads for each strain were therefore pooled for all subsequent analysis. For full statistics, see **Table S1**.

### Sequencing yield

During the 48-hour MinION sequencing run, 1,803,648 reads were generated, equating to 9.73 Gbp of sequence. 28.78 % of these reads (574,053 reads, 2.8 Gbp) were not assigned to the correct barcode bin during demultiplexing, leaving 6.93 Gbp (1,229,595 reads) of useable sequencing data (**Fig. 1b**). Normalised yield per barcode (taking into account ng of gDNA included in the pooled sequencing library) was particularly high for one barcode (NB11, 15.28 Mbp ng^-1^) but otherwise relatively consistent, ranging from 7.38 to 10.28 Mbp ng^-1^ with a mean yield of 9.06 Mbp ng^-1^ (std. error=0.37). Mean read length for the full read set was 5,689 bp. For full statistics including which barcode was assigned to each sample, see **Table S1**.

### Assembly tool testing – nanopore-only

**Table 1** shows the quality measurements for the best nanopore-only assembly per tool trialled. All tools tested were able to resolve the nanopore long reads for UK36 into a complete, closed contig, using default assembly options with no manual intervention. In total, 97 different tool combinations were trialled. Alignment of drafts from different tools using progressiveMauve revealed that each tool also assembled the genome into the same arrangement (**Fig. S1**). However, the length of the draft assemblies showed greater variation: 3.984 to 4.134 Mbp, with a mean length of 4.108 Mbp.

**Table 1:** best *de novo* assembly options and quality measurements for nanopore-only assemblies

| Assembler | Pre-assembly read correction | Pre-assembly read filtering / x coverage | Rounds of Racon polishing | Polishing with Nanopolish | Contigs | Assembly length / Mbp | % identity compared to Illumina-only | BUSCOs present/fragment/missing (of 40) |
| --- | --- | --- | --- | --- | --- | --- | --- | --- |
| ABRuijn | Yes | No | 0 | Yes | 1 | 4.105 | 99.59 | 37/2/1 |
| Canu | Yes | No | 4 | Yes | 1 | 4.133 | 99.54 | 36/1/3 |
| Flye | Yes | No | 0 | Yes | 1 | 4.108 | 99.56 | 35/3/2 |
| Miniasm + Minimap | Yes | No | 5 | Yes | 1 | 4.111 | 99.55 | 37/0/3 |
| Unicycler | Yes | No | 8* | Yes | 1 | 4.107 | 99.55 | 35/2/3 |

\*The rounds of Racon listed for Unicycler were carried out as part of the Unicycler protocol; no manual rounds of Racon were conducted

Comparing like-for-like assemblies before and after polishing shows that Nanopolish improves identity by 0.216 % on average (n=16, paired t-test p<0.001). Polishing with Racon produced inconsistent results: the identity decreased after Racon polishing of ABruijn and Flye drafts, increased by 2.01 % after the optimal number of polishing rounds for pre-corrected non-ABruijn/Flye drafts (n=3), and increased by 15.15 % after optimal rounds for non-ABruijn/Flye drafts with no pre-correction (n=4). The mean number of Racon polishes required to reach optimal % identity (after which % identity began to decrease) was 4.75 (n=7).

The assembly with greatest % identity compared to the Illumina-only draft (99.59 %) combined pre-assembly read correction with Canu, assembly with ABruijn and post-assembly polishing with Nanopolish. The assemblies were also assessed using BUSCO, which searches draft assemblies for copies of Benchmarking Universal Single-Copy Orthologs (BUSCOs). BUSCOs are sets of core genes which are likely to appear universally in related organisms. A set of 40 such core genes from the *Escherichia coli* genome are used as the gram-negative bacterial BUSCOs; if a genome has been assembled accurately, BUSCO is more likely to be able to identify these 40 genes within its sequence. Of the drafts assessed here, the ABruijn assembly contained the highest number of identifiable BUSCOs (37 full and 2 partial, of the full set of 40).

### Assembly tool testing – hybrid

A number of hybrid assembly strategies were trialled, including polishing a long-read assembly with short reads, scaffolding short-read contigs with long reads, and using both short and long reads together during assembly (Table 2 shows the best draft produced by each tool). Scaffolding short-read contigs with long reads using SPAdes produced one of the highest accuracy assemblies (99.68%), but did not fully resolve the genome, as six contigs remained. No further polishing was attempted with this SPAdes assembly, as polishing would not close the remaining gaps between the contigs.

**Table 2:** best *de novo* assembly options and quality measurements for hybrid assemblies

| Assembler | Pre-assembly read correction | Pre-assembly read filtering / x coverage | Assembly includes short reads? | Rounds of Racon polishing | Polishing with Nanopolish | Rounds of Pilon polishing | Contigs | Assembly length / Mbp | % identity compared to Illumina-only | BUSCOs present/fragment/missing (of 40) |
| --- | --- | --- | --- | --- | --- | --- | --- | --- | --- | --- |
| ABRuijn | Yes | No | No | 0 | Yes | 3 | 1 | 4.109 | 99.67 | 40/0/0 |
| Canu | Yes | No | No | 4 | Yes | 3 | 1 | 4.130 | 99.66 | 40/0/0 |
| Flye | Yes | No | No | 0 | Yes | 3 | 1 | 4.108 | 99.67 | 40/0/0 |
| Miniasm + Minimap | No | No | No | 5 | Yes | 4 | 1 | 4.107 | 99.66 | 40/0/0 |
| SPAdes | Yes | No | Yes | n/a | n/a | n/a | 6 | 4.105 | 99.68 | 40/0/0 |
| Unicycler | Yes | No | Yes | 4* | No | 8* | 1 | 4.107 | 99.68 | 40/0/0 |

\*The rounds of Racon and Pilon listed for Unicycler were carried out as part of the Unicycler protocol; no manual rounds of polishing were conducted for this assembly

The best hybrid assemblies per tool were significantly more accurate than the best nanopore-only assemblies per tool, with a mean identity improvement of 0.11 % (hybrid n=6, nanopore-only n=5, paired t-test p<0.001). In addition, all hybrids contained all 40 identifiable BUSCOs, and all non-SPAdes hybrids were single closed contigs and showed the same arrangement when aligned using progressiveMauve (fig S2).

The best single-contig hybrid assembly, with 99.68 % identity, was produced using Unicycler’s hybrid option. Table S2 shows the results from all nanopore-only and hybrid tests.

### Assembly & alignment of all strains

Using the nanopore-only and hybrid pipeline developed through the tests described here (**Fig. 2**), draft genomes were assembled for all five UK strains sequenced during our barcoded run. The assemblies were assessed for % identity compared to each strain’s Illumina-only assembly, GC content, genome length and number of key IS element features; they were also annotated using Prokka. The full results of this analysis are shown in **Table 3**.

**Table 3:** Assembly statistics for five UK *B. pertussis* strains, assembled using our hybrid pipeline

| Pipeline | Strain | Contigs | Genome length / Mbp | GC content / % | % identity compared to Illumina-only | # genes predicted | IS481 copies | IS1002 copies | IS1663 copies |
| --- | --- | --- | --- | --- | --- | --- | --- | --- | --- |
| NANOPORE-ONLY | UK36 | 1 | 4.108 | 67.69 | 99.47 | 4698 | 258 | 8 | 17 |
|  | UK38 | 1 | 4.108 | 67.69 | 99.49 | 4741 | 258 | 8 | 17 |
|  | UK39 | 1 | 4.109 | 67.70 | 99.48 | 4588 | 258 | 8 | 17 |
|  | UK48 | 1 | 4.114 | 67.70 | 99.47 | 4610 | 262 | 8 | 17 |
|  | UK76 | 1 | 4.113 | 67.70 | 99.32 | 4608 | 262 | 8 | 17 |
| HYBRID | UK36 | 1 | 4.107 | 67.70 | 99.68 | 3757 | 258 | 8 | 17 |
|  | UK38 | 1 | 4.108 | 67.70 | 99.69 | 3757 | 258 | 8 | 17 |
|  | UK39 | 1 | 4.108 | 67.70 | 99.69 | 3804 | 258 | 8 | 17 |
|  | UK48 | 2 | 4.112 | 67.70 | 99.68 | 3763 | 262 | 8 | 17 |
|  | UK76 | 1 | 4.113 | 67.70 | 99.54 | 3753 | 262 | 8 | 17 |

**Fig. 2:**
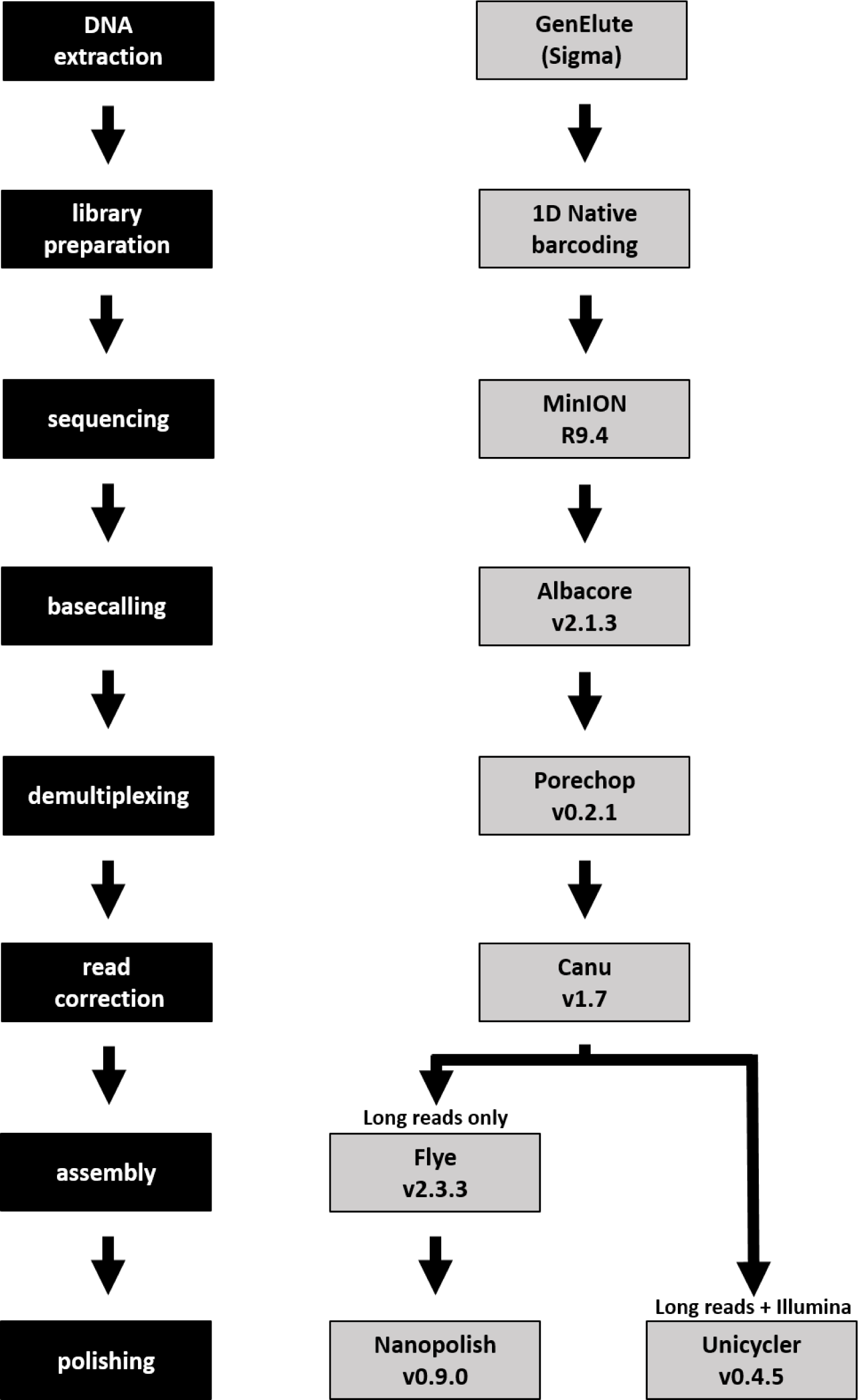
Our nanopore-only and hybrid sequencing pipelines, developed through extensive testing of available tools.

In the hybrid assemblies, two strains, UK48 and UK76, had longer genomes than the others (4.112 and 4.113 Mbp respectively, compared to 4.108 Mbp), which corresponds with them also having more copies of the most abundant IS element, IS481. All strains but one were assembled into single contigs. The remaining strain, UK48, was assembled into five contigs (N50=3.934 Mbp). Of these, three were shorter than 500 bp, and were subsequently discarded. The remaining two contigs were 3,934,355 bp and 178,023 bp. Mapping the raw UK48 reads to the Tohama I reference sequence revealed a section of around 200 kbp, located between 1.4 and 1.6 Mbp, which had double the read depth of the rest of the reference; the doubled read depth suggests that this section of the genome is duplicated in UK48. No other strain had a similarly duplicated section, although the coverage of UK76 was enriched by around 25 % at the same locus (**Fig. 3**). These abnormalities are also present in the Illumina reads, which were sequenced approximately five years before our nanopore reads (**Fig. S3**).

**Fig. 3:**
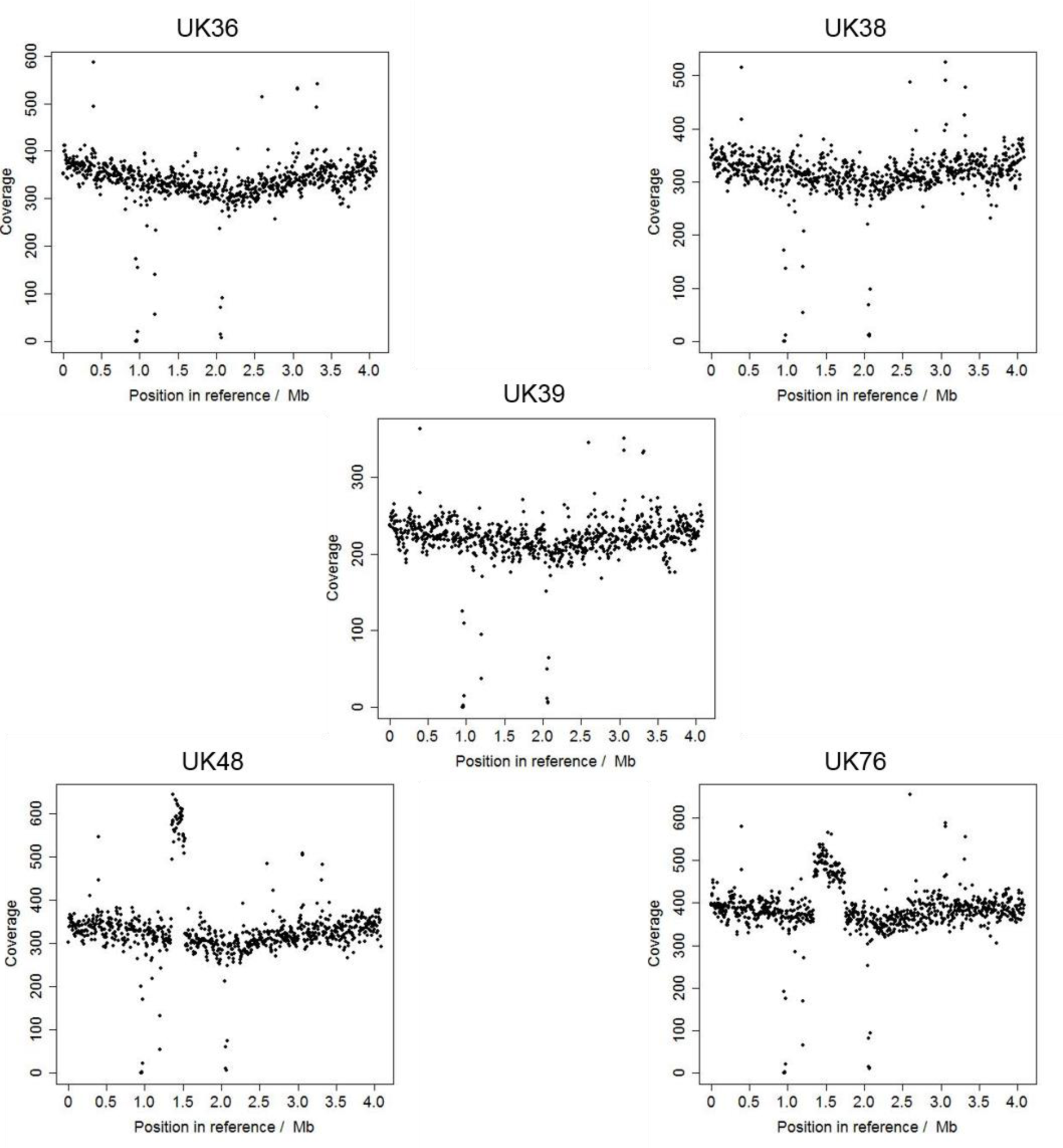
Raw read coverage of the Tohama I reference genome for each of the UK strains sequenced. Raw reads(Albacore and Porechop) were aligned to the Tohama I reference sequence using BWA MEM, and coverage assessed by Samtools depth. The coverage of three strains (UK36, UK38 and UK39) was consistent across the whole reference genome, whereas UK48 and UK76 coverage was enriched in certain locations. In UK48, a large section around 1.4 to 1.6 Mbp into the reference appears to have exactly twice as much coverage as the rest of the genome. In UK76, a section from 1.4 to 1.7 Mbp is enriched by one quarter. In addition, there are sections of low coverage at 1 Mbp and 2Mbp in every strain sequenced here; these likely correspond to parts of the reference genome which have been lost since 2003, or which the UK strains never possessed.

The hybrid assembly for one strain, UK76, had slightly lower % identity (99.54 %) than the other strains, each compared to their respective Illumina-only ABySS assembly. Discounting UK76, the assemblies had a mean identity of 99.69 % (n=4). The GC content of the strains varied little: the content for all strains was 67.70 % when rounded to 2 d.p. The number of genes predicted by Prokka was also relatively consistent, varying from 3757 to 3804.

The UK36 proteins predicted by Prokka were assessed by Ideel, which searched the Trembl database [45] for similar proteins. The length of the Prokka-predicted proteins was divided by those of the identified similar Trembl proteins; a perfect match would equal 1.0. This method, therefore, indicates whether indels in a draft sequence cause frameshifts which subsequently lead to truncated (or over-long) protein prediction. After manual curation to remove known real pseudogenes, over 98% of Prokka-predicted genes had a Prokka:Trembl length ratio of greater than 0.9. This suggests that the residual error in the hybrid assemblies does not cause substantial annotation problems, so the hybrid assemblies for all five strains were submitted to GenBank (accession numbers CP031289, CP031112, CP031113, QRAX00000000, CP031114).

All strains were assembled into single contigs using the nanopore-only pipeline. These assemblies were aligned using progressiveMauve (Fig. 4), displaying extensive genomic rearrangement between strains; three, UK36, UK38 and UK39, shared exactly the same arrangement, whilst UK48 and UK76 were rearranged.

**Fig. 4:**
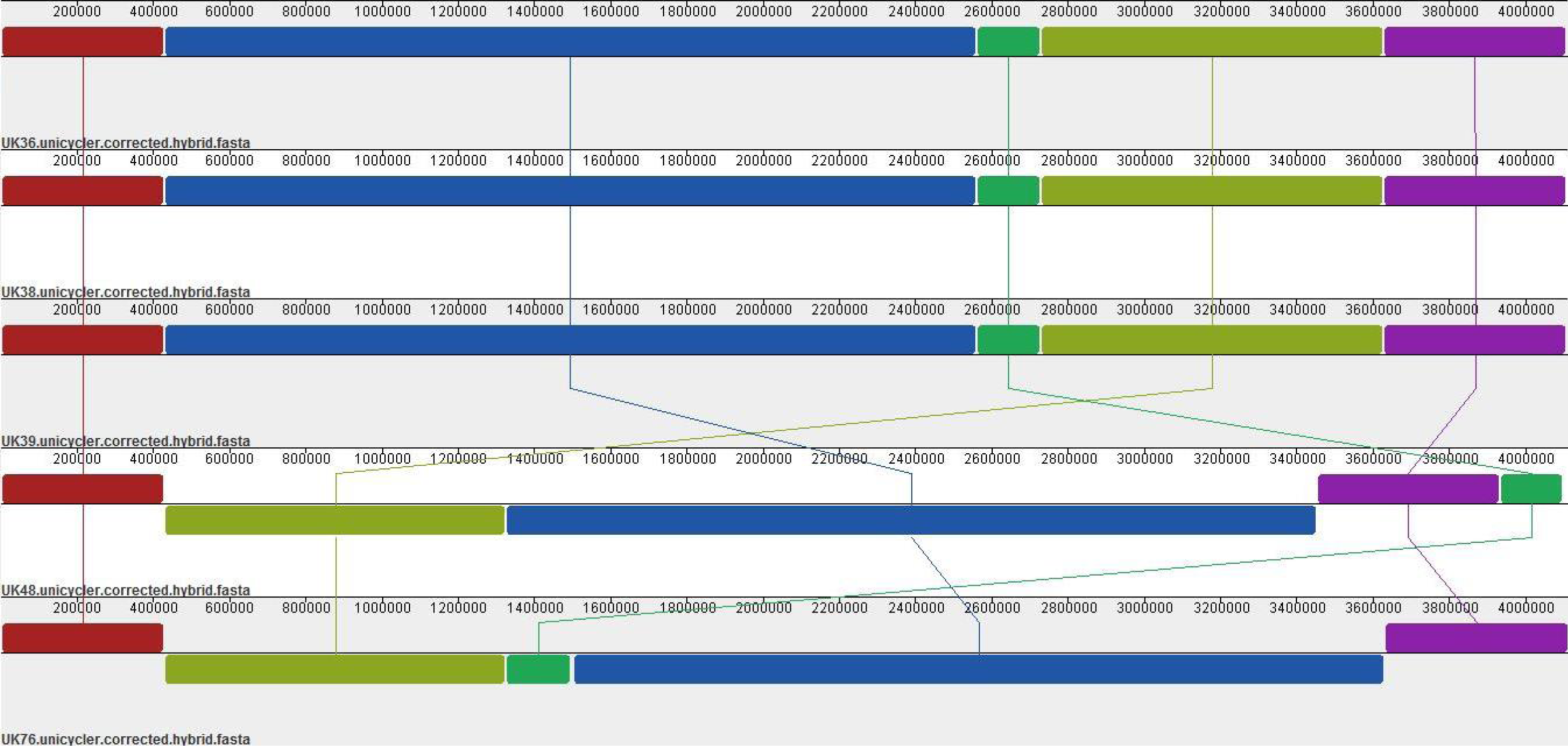
Alignment of our five sequenced strains, showing genomic rearrangement. Our five UK *B. pertussis* strains (UK36, UK38, UK39, UK48 and UK76) were assembled using our nanopore-only pipeline, resulting in single, closed-contig, assemblies. The closed assemblies were aligned with progressiveMauve, which showed that even strains which are closely temporally related can display different genomic arrangements.

## DISCUSSION

### Accuracy of long-read sequencing is improving but error estimation is challenging

Our primary aim in this study was to determine whether long reads produced by nanopore sequencing using ONT’s MinION can be used to produce closed *B. pertussis* genome sequences by *de novo* assembly. In addition, we trialled numerous data analysis strategies - including different basecallers, different assembly and polishing tools, and long-read-only vs hybrid assembly - to determine the optimal analysis pipeline which can currently produce the most accurate genome sequences.

Until mid-2017, the only nanopore basecalling option was MinKNOW, the software provided with the MinION. Albacore and several other stand-alone basecalling tools are now available and appear to offer improved basecalling accuracy [39]. We selected Albacore to compare with MinKNOW due to its ease of use and demultiplexing ability. The read sets from Albacore were clearly superior; a 2.46 % mean accuracy improvement across 9.73 Gbp of raw sequencing reads equates to 239 million base errors corrected. We also saw a 0.088 % improvement in each 4.1 Mbp Albacore assembly compared to the equivalent MinKNOW assemblies, equating to over 3,500 fewer errors per genome.

Without a recent, closely-related reference sequence, error estimation in *B. pertussis* assemblies is inexact. Comparison with the 2003 Tohama I reference sequence will identify basecalling errors which are false positives, having arisen due to natural variation between different strains (that is, true SNPs will be identified as errors). Moreover, the validity of Tohama I as a representative of all *B. pertussis* strains is questionable [46]. The Illumina reads available for four of our sequenced strains (UK 36, 38, 39 and 48) showed 98.44 % identity with the Tohama I sequence, suggesting natural genetic variation between Tohama I and these UK strains of around 1.5 %. The false positive rate is thus around 1.5 % when using Tohama I to assess assembly accuracy. On the other hand, comparison with Illumina-only assemblies requires short read data to be available, and assumes the Illumina reads to be close to 100 % accurate, which could be a flawed assumption. The Illumina reads for UK76, for example, had raw identity of only 87.32 % compared to Tohama I. With no distinctive features noted for UK76 in our assembly or in the original comparison of UK epidemic strains [7], it is unlikely that the UK76 genome is really 11 % less like Tohama I than the other strains sequenced here. It seems more likely that the Illumina reads are inaccurate; if this is the case, our assessments of the accuracy of our UK76 assemblies were skewed. This could explain why our UK76 hybrid assembly had a slightly lower estimated accuracy than the other strains. Compared to Tohama I, our hybrid UK76 assembly showed 98.49 %, similar to the identity of our other hybrid assemblies (n=5, mean=98.57 %), suggesting that the inaccuracies of the raw Illumina reads do not translate into inaccuracies in the final assembly; only our estimation of accuracy by comparison to the Illumina-only draft is affected. Overall, neither comparison to the Tohama I reference nor comparison to an Illumina-only assembly is ideal for assessing error when working with novel strains, and neither strategy gives us a completely accurate estimate, but using a combination of both comparisons allows a good estimate of assembly error.

Having estimated our hybrid assemblies to be, on average, 99.69 % accurate, we can conclude that roughly 13,000 bases in each 4.1 Mbp draft genome are incorrect. Whilst these incorrectly called bases will not influence comparisons of genome arrangement (as shown in **Figs. S1 and S2**), residual base errors in draft genome sequences assembled using long reads remain a concern, with the potential to falsely identify SNPs or prevent accurate protein prediction [47]. Incorrect sequencing of homopolymers is a known weakness of many sequencing methods, including nanopore sequencing [17], and our assemblies are no exception. Indeed, a base-level manual comparison of one of our hybrid assemblies with a more accurate Illumina-only draft using progressiveMauve revealed that every difference occurred in a homopolymeric tract, with the hybrid sequence having inserted or deleted bases. Two options for correct SNP identification, therefore, are manual correction of known homopolymeric indels [47], and simply ignoring SNPs which appear to occur in homopolymeric regions. The manual correction option would be time-consuming, whilst the second option could result in false negatives. Nevertheless, until improved pore chemistry or basecalling tools are available which do not produce homopolymeric indels, the use of either option means that SNP identification is still possible, even in assemblies which are less than 100 % accurate.

Correct prediction of proteins appears to be of less concern than SNP identification in our hybrid assemblies: all 40 potential bacterial BUSCOs were present in full for all of our strains, and both Quast and Prokka were able to identify the majority of the Tohama I reference proteins in the same assemblies. In addition, assessment of our UK36 hybrid using Watson’s Ideel pipeline [42] suggested that, although we know some errors remain, they do not substantially inhibit the correct prediction of full-length proteins during annotation. It is here, however, that we can clearly see the benefit of the hybrid assemblies over the nanopore-only assemblies: although the mean accuracy of the nanopore-only assemblies (99.48 %) was only 0.2 % lower than that of the hybrids, none of the strains contained full copies of all 40 BUSCOs.

### Does the de Bruijn graph method assemble highly repetitive prokaryotic genomes more accurately than other commonly used methods?

The opinion of the sequencing community has long been that de Bruijn graph assembly is not as effective for error-prone long reads as other *de novo* assembly methods [48, 49]. The tool which consistently produced the most accurate nanopore-only *B. pertussis* assemblies was therefore unexpected: the % identity and indel rates of our ABruijn assemblies were better by far than those of the Canu, Miniasm or Unicycler assemblies. The recent version change of ABruijn to Flye seems to have negatively affected these metrics in some of our strains; however, whilst the ABruijn assemblies were better than the Flye assemblies, the Flye assemblies were still better than those produced by other tools. Another recent study, which assembled highly complex and repetitive *Pseudomonas koreensis* genomes using ultra-long nanopore reads, also found Flye to produce the most accurate assemblies [16]. This suggests that the de Bruijn method might be optimal for prokaryotic genomes which contain a high number of repeats.

### Are residual unresolved ultra-long repeats present in some strains?

The region of enriched coverage between 1 and 2 Mbp in the Tohama I reference genome observed in the UK48 reads (**Fig. 3**) is likely to indicate a large (around 200 kbp) duplication of that region which is present in UK48 but not in the reference. A less obvious duplication may also be present in the genome of UK76: a 400 kbp region between 1 and 2 Mbp shows 125 % coverage. An alternative potential cause for these coverage abnormalities is contamination of the sequencing library, particularly from UK48 into UK76. However, the presence of the same abnormalities in other read sets for both strains suggests that they have not been caused by such contamination (**Fig. S3**). Neither Flye nor Unicycler however, was able to resolve the duplication correctly. Our UK48 reads had a mean length of 6,243 bp, whilst the UK76 read mean length was 5,480 bp; if the key to resolving long repeats is to use reads longer than the longest repeat, we will need ultra-long reads in the order of hundreds of thousands of bases to resolve these putative duplications [16, 17]. Methods to extract and sequence such long reads have been developed by the nanopore community, with reports of reads in the order of millions of bases [50].

### Two possible pipelines for *B. pertussis* genome sequence resolution

We have shown here that resolution of five *B. pertussis* genomes per MinION flow cell is possible, whether using long reads alone, or in combination with short reads. We used ten barcoded samples but were able to pool pairs of samples because reads from FFPE-repaired and non-FFPE-repaired libraries were of comparable quality. Consequently, around one fifth of all usable reads belonged to each strain, equating to over 300x *B. pertussis* genome coverage per strain. 300x coverage probably exceeds that required to achieve comparable results: a draft produced from just the non-FFPE-repaired reads for UK36, pre-corrected and assembled with Flye, had an identity of 99.467 %, whilst the same assembly produced by the pooled FFPE-repaired and non-FFPE-repaired reads had an identity of 99.474 %. The non-FFPE-repaired reads alone had 175x coverage, less than half the 359x coverage of the full set of pooled reads. This suggests that twice as many strains could be *de novo* assembled per flow cell without a notable drop in accuracy. Thus, resolution of ten *B. pertussis* genomes per MinION flow cell should be possible.

If short reads are also available, we have shown that hybrid assembly, using pre-correction with Canu followed by Unicycler, remains the most accurate method. Indeed, for now, for full strain characterisation (including comparison of genome arrangement, SNP identification and allele-typing), hybrid assemblies are required. For comparison of genome structure and arrangement only (e.g. **Fig. 3**), however, our nanopore-only pipeline, which uses Canu pre-correction, Flye assembly and post-assembly polishing with Nanopolish, can produce single contig assemblies of adequate accuracy for all but the most unusual *B. pertussis* genomes.

### Continued improvement of long-read data processing tools

Although the pipelines we have developed here produce the most accurate *B. pertussis* genome sequences currently possible, the tools available for the analysis of nanopore sequencing data are continually improving. A recent update to Racon added the ability to polish assemblies with Illumina reads; a brief comparison of this with Pilon, however, showed little improvement to our data, so we did not add short-read Racon polishing to our suite of tests. For basecalling, we chose to compare MinKNOW with Albacore, but other basecalling tools are already available, and more still are under current development [39]. Alternative basecallers such as Chiron [51] or the currently in-development Guppy, which use entirely new basecalling algorithms, may offer further accuracy improvements and could be trialled with existing and future *B. pertussis* data sets, particularly if Illumina short reads are not available for hybrid assembly.

We tested the most commonly used *de novo* assembly tools suitable for long reads and, at the time of writing, are not aware of any newly-released tools. However, minor (or sometimes major, in the case of ABruijn to Flye) updates are common. New polishing tools are also being developed: ONT’s own Medaka, for example, is claimed to rival Nanopolish in terms of speed and assembly improvement capabilities [52]. In addition, MaSuRCA [53] was not trialled here due to the low Illumina coverage (the manual suggests 50x+ for hybrid assemblies, whereas we had only 37.5x coverage for UK36). Ultimately, for the foreseeable future, no data pipeline including nanopore reads should be set in stone; we will continue to trial new tools and to update our pipeline where appropriate, and would suggest that similar pipeline optimisation may be required for each organism to be sequenced.

## Acknowledgements

We are grateful to Public Health England’s National Reference Laboratory for providing the strains sequenced here. Many thanks are also due to Dan Turner, Dan Fordham and Phill James for hosting NR at Oxford Nanopore Technologies to conduct the barcoded nanopore sequencing. The data analysis performed here would not have been possible without access to the fantastic bioinformatics resource, CLIMB (developed by the MRC, grant number MR/L015080/1). Finally, we would like to acknowledge Ryan Wick, who unknowingly inspired the highest level of open-accessibility to which we have tried to aspire here.

## ABBREVIATIONS

BUSCO: Benchmarking Universal Single-Copy Orthologs
CLIMB: Cloud Infrastructure for Microbial Bioinformatics
FFPE: Formalin-Fixed, Paraffin-Embedded
FHA: filamentous haemagglutinin
IS: insertion sequence
MRC: Medical Research Council
ONT: Oxford Nanopore Technologies
Prn: Pertactin
Pt: Pertussis toxin
SPRI: Solid Phase Reversible Immobilization

